# Cerebral malaria is regulated by host mediated changes in *Plasmodium* gene expression

**DOI:** 10.1101/2022.11.23.517617

**Authors:** Clare K. Cimperman, Mirna Pena, Sohret M. Gokcek, Brandon P. Theall, Meha V. Patel, Anisha Sharma, ChenFeng Qi, Daniel Sturdevant, Louis H. Miller, Patrick L Collins, Susan K. Pierce, Munir Akkaya

**Author notes:** Address correspondence to Munir Akkaya.

## Abstract

Cerebral Malaria (CM), the deadliest complication of *Plasmodium* infection, is a complex and unpredictable disease. However, our understanding of the host and parasite factors that cause CM is limited. Using a mouse model of CM, experimental CM (ECM), we performed a three-way comparison between: ECM susceptible C57BL/6 mice infected with ECM-causing *Plasmodium* ANKA (*Pb* ANKA) parasites (ANKA^(C57BL/6)^); ECM resistant Balb/c mice infected with *Pb* ANKA (ANKA^(Balb/c)^); and C57BL/6 mice infected with *Pb* NK65 that does not cause ECM (NK65^(C57BL/6)^). All ANKA^(C57BL/6)^ mice developed CM. In contrast in ANKA^(Balb/c)^ and NK65^(C57BL/6)^ infections do not result in CM and proceed similarly in terms of parasite growth, disease course and host immune response.. However, parasite gene expression in (ANKA^(C57BL/6)^) was remarkably different than in ANKA^(Balb/c)^ but similar to the gene expression in NK65^(C57BL/6)^. Thus, *Pb* ANKA has a ECM-specific gene expression profile that is only activated in susceptible hosts providing evidence that the host has a critical influence on the outcome of infection.

## Introduction

Cerebral malaria (CM) is the deadliest complication of *Plasmodium* infection^1^. Despite occurring in only approximately 1% of infected individuals, CM is responsible for over 90% of all malaria parasite related deaths which equals to around 400,000 fatalities each year^2, 3^. According to WHO, CM affects mostly young African children and is estimated to be 15-25% fatal even with effective anti-malarial drug treatment ^3, 4^. Moreover, survivors of the disease often suffer from long term neurological sequela dramatically decreasing their quality of life^5, 6^. CM is a complex disease characterized by sequestration of infected red blood cells (iRBCs) in brain vasculature; migration of innate and adaptive immune cells to brain parenchyma; increased production of proinflammatory mediators altogether leading to multifocal intracranial hemorrhage and brain edema^3, 7-11^. Studies on the pathophysiology of CM are limited due to both rapid progression of disease -often fatal within 48 hours after the onset of neurological symptoms- and limited access to pre- and post-mortem patient material^12-14^. Therefore, we have a limited understanding of the pathogenesis of CM and, why *Plasmodium* infection in some patients progress into CM, while in vast majority of patients, the disease runs a course free of any neurological symptoms is still unclear.

Studies of CM in mice, known as experimental CM (ECM) have been used extensively to delineate the pathogenesis of CM^15, 16^. In particular, *Plasmodium berghei* (*Pb*) ANKA infection of C57BL/6 host has been shown to recapitulate the majority of the CM symptoms^17-21^. Further research using *Pb* ANKA parasites in different hosts showed that although most mouse strains can be infected by *Pb* ANKA only certain strains such as C57BL/6 and CBA/Ca were susceptible to ECM, whereas other strains such as Balb/c and DBA/2J were resistant ^22-25^. These findings pointed out the importance the of host in determining the outcome of parasite infection. In separate studies, infection of the ECM susceptible C57BL/6 strain with a different *Plasmodium berghei* strain (*Pb* NK65) that is closely related to *Pb* ANKA had no ECM phenotype^26^. Genomic analyses showed that *Pb* ANKA and *Pb* NK65 parasites differed by only 20 single nucleotide polymorphisms (SNPs) in their coding regions^26, 27^. Recently, using Crispr-Cas9 mediated point mutation in one of these SNPs located in the DNA binding domain of an ApiAP2 transcription factor, we reported major changes in the development of host immune response against the parasite and the outcome of parasite infection albeit no change in ECM phenotype^26, 28^. Altogether, these reports highlighted the importance of parasite genetic factors in determining the outcome of the disease.

Although, such studies established that both host and parasite related factors determine whether or not *Plasmodium* infections lead to ECM, research aimed to identify how host and parasite factors influence each other is limited. Clearly a better understanding of the contribution of parasite and host factors in mouse models would benefit efforts to determine the how human to human variation as well as continuously changing parasite gene expression determines the outcome of human *Plasmodium* infection.

Here we performed a three-way comparison between ECM causing *Pb* ANKA in ECM susceptible C57BL/6 host (ANKA^(C57BL/6)^); ECM causing *Pb* ANKA in ECM resistant Balb/c host (ANKA^(Balb/c)^); and non-ECM causing *Pb* NK65 in ECM susceptible C57BL/6 host (NK65^(C57BL/6)^). Our comprehensive analyses showed that host influences the disease progression by modulating parasite gene expression.

## Results

### ANKA^(Balb/c)^ and NK65^(C57BL/6)^ infections have similar disease courses

We monitored the progression of malaria following ANKA^(C57BL/6)^, ANKA^(Balb/c)^ and NK65^(C57BL/6)^ *Plasmodium* infection conditions (Supplemental Fig. 1a). Only ANKA^(C57BL/6)^ rapidly progressed to severe disease as evidenced by early uniform mortality with all mice dying by day 7 p.i. (Fig.1a). Histopathological evaluations of brains taken from a group of animals on day 6 p.i. proved ECM phenotype in ANKA^(C57BL/6)^ group which showed multifocal hemorrhages (Fig 1b, c). These hemorrhages localized particularly in cerebellum (Fig.1b) and olfactory bulb (Fig.1c) which were formerly demonstrated as areas that are preferentially affected by ECM^16, 26, 29^. On the other hand, ANKA^(Balb/c)^ and NK65^(C57BL/6)^ groups both had longer disease courses (Fig.1a) and no brain pathology (Fig 1b,c) confirming that these two conditions do not develop ECM.

**Figure 1.**
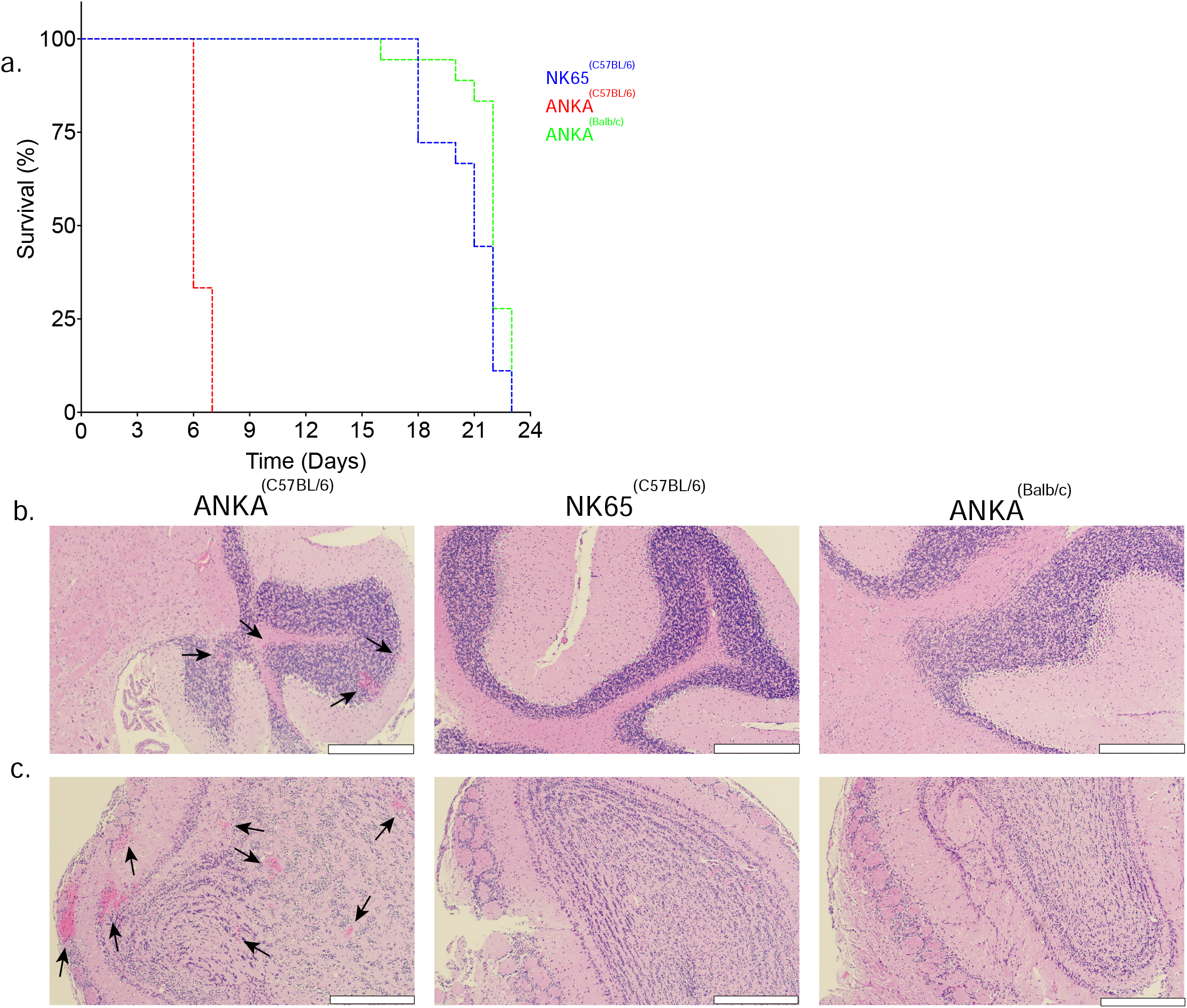
ANKA^(Balb/c)^ and NK65^(C57BL/6)^ infections have similar disease courses that is distinct from ANKA^(C57BL/6)^. Mice were infected with 10^6^ iRBCs from relevant parasites. **a)** Kaplan Meier graph showing the survival percentage for each group through the course of the disease. Each group contains 20 infected mice. **b**,**c)** Hematoxylin & Eosin stained brain tissue sections taken at day 6 post infection. Close up images of cerebellum **(b)** and olfactory bulb **(c)** are shown. Arrow heads point sites of local hemorrhage. White scale bars on lower left corner represent 200 μm. Images represent three brains per group.

We further characterized the progression of blood stage malaria in ANKA^(C57BL/6)^, ANKA^(Balb/c)^ and NK65^(C57BL/6)^ via regular blood checks for progression of parasitemia and hemoglobin levels. *Pb* ANKA infected mice (ANKA^(Balb/c)^ and ANKA^(C57BL/6)^) regardless of their host showed a faster increase in parasitemia as compared to NK65^(C57BL/6)^ within the first few days p.i. (Fig. 2a) which may suggest an inherent proliferation advantage for *Pb* ANKA as compared to *Pb* NK65. However, due to ECM mediated mortality by day 7 p.i. no further measurements could be made from ANKA^(C57BL/6)^. Later in the infection around 6 days pi parasitemias continued to increase with ANKA^(Balb/c)^ reaching higher levels of parasitemia (60%) as compared to NK65^(C57BL/6)^ (30%). The higher levels of parasitemia in ANKA^(Balb/c)^ as compared to NK65^(C57BL/6)^ may contribute to the slightly higher clinical scores in ANKA^(Balb/c)^ parasite despite the lack of ECM phenotype. Reduction in hemoglobin levels were comparable between ANKA^(Balb/c)^ and NK65 ^(C57BL/6)^ especially in later stages of the disease despite higher starting hemoglobin levels in Balb/c mice prior to infection. Progressive decrease in hemoglobin levels ANKA^(Balb/c)^ and NK65 ^(C57BL/6)^ is in line with previous observations which report anemia as the cause of death in these non-ECM cases^28, 30-33^.

**Figure 2.**
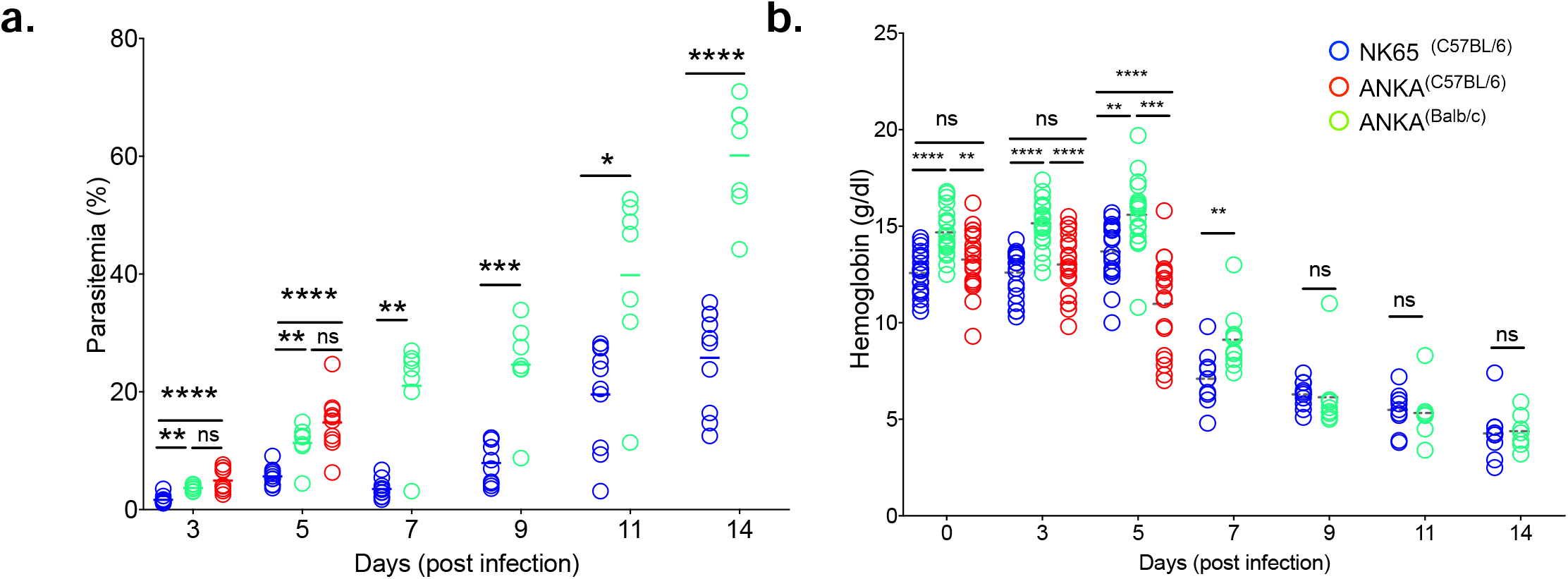
ANKA^(Balb/c)^ and NK65^(C57BL/6)^ infections cause mortality due to anemia. **a**,**b)** Mice were infected as outlined in Figure 1. Changes in parasitemia (percentage of infected RBC among total RBC) **(a)** and hemoglobin concentration **(b)** are measured through routine blood checks. Each circle represents an individual animal. (ns = P > 0.05; * = 0.01 < P ≤ 0.05; ** = 0.001 < P ≤ 0.01; *** = 0.0001 < P ≤ 0.001. (Comparisons between three groups were made using One Way ANOVA with Tukey test. Comparisons between two groups were made using Welch’s t test) Data represents two experiments.

### ANKA^(Balb/c)^ and NK65^(C57BL/6)^ infections trigger a stronger immune response as compared to ANKA^(C57BL/6)^ infection

Having observed differences in development of disease symptoms and progression of parasitemias (Fig 1 and Fig 2), suggest the possibility that ANKA^(C57BL/6)^, ANKA^(Balb/c)^,and NK65^(C57BL/6)^ infections trigger host immune response differently. We analyzed spleen samples collected on day 6 pi and characterized the changes in immune cell numbers and their activation/differentiation states (Fig 3). ANKA^(Balb/c)^, and NK65^(C57BL/6)^ infected groups had similar numbers of total B cells (Fig.3a), CD^4+^ T cells (Fig.3b) and CD8^+^ T cells (Fig.3c) in their spleens. In contrast, ANKA^(C57BL/6)^ had significantly lower numbers of all three cell types indicating differences in immune responses. We analyzed B cell differentiation in terms of the expansion of cells in the germinal center (GC) and plasma cell (PC) lineages (Fig.3d-g). Absolute numbers of GC and PC lineage cells were similar for ANKA^(Balb/c)^ and NK65^(C57BL/6)^ which were uniformly higher as compared to ANKA^(C57BL/6)^ (Fig.3d, f). In addition to higher absolute numbers for GC and PC lineage cells for ANKA^(Balb/c)^,and NK65^(C57BL/6)^, the percentage of GC and PC lineage cells within the parent population, B cells and live splenocytes respectively, also increased as compared to ANKA^(C57BL/6)^ condition (Fig.3e, g). These observations rule out that increases in absolute numbers of GC and PC are merely a reflection of the overall increase in total B cell numbers rather than B cell differentiation. In line with these observations in B cell differentiation, in ANKA^(Balb/c)^, and NK65^(C57BL/6)^ infections as compared to ANKA^(C57BL/6)^ infections, follicular helper T cells (T_fh_) and activated CD8^+^ T cells were higher in both as absolute numbers and as percentages of total CD4^+^ and CD8^+^ T cells respectively (Fig.3h-k). Lastly, immunohistochemistry analysis of spleen sections showed, in addition to the lower GC numbers in ANKA^(C57BL/6)^ (Fig.3d, e), GC cells failed to organize in proper clusters (Fig.3l). Altogether our findings show an inefficient and low-grade immune response against ECM causing ANKA^(C57BL/6)^ as compared to non-ECM causing groups.

**Figure 3.**
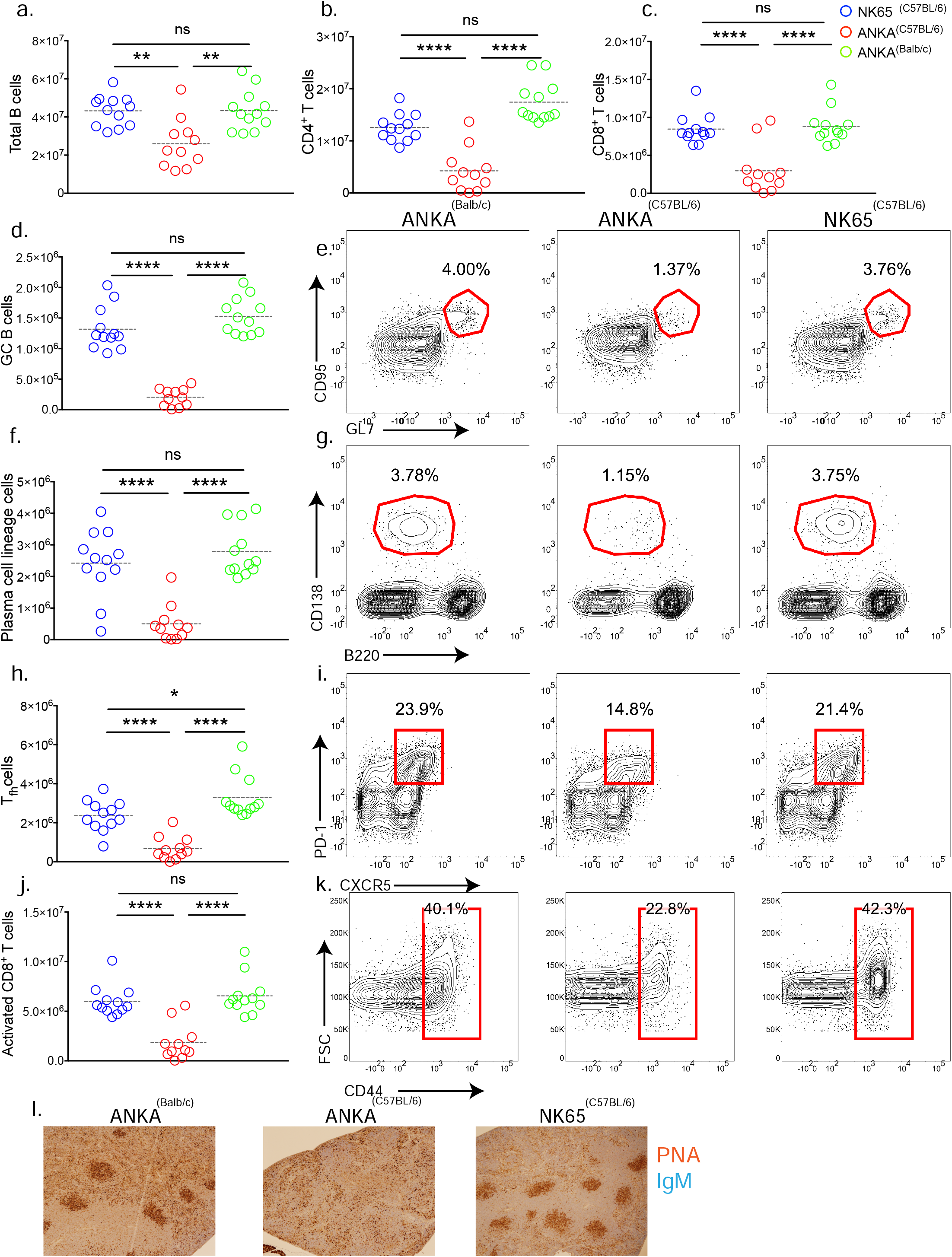
ANKA^(Balb/c)^ and NK65^(C57BL/6)^ infections induce robust immune response. **a-l)** Mice were infected with parasites as outlined in Figure 1. Spleens were harvested on day 6 post infection and development of immune response was characterized in flow cytometry **(a-k)** or using immunohistochemistry **(l)**. Graphs show absolute numbers of each cell group. Circles represent individual mice and dotted lines refer to means **(a-d, f, h, j)**. Representative flow cytometry contour plots showing the frequencies of GCs among B cells **(e)**, plasma cells among viable splenocytes **(g)**, T_fh_ cells among CD4^+^ T cells **(i)** and activated CD8^+^ T cells among total CD8^+^ T cells **(k)** are shown. **l)** Spleen sections stained with PNA and IgM to visualize localization and architecture of GCs in infected mice. (ns = P > 0.05; * = 0.01 < P ≤ 0.05; ** = 0.001 < P ≤ 0.01; *** = 0.0001 < P ≤ 0.001. One Way ANOVA with Tukey test). Experiments were repeated three times.

### Host has a strong influence on parasite gene expression

Thus far our data showed that ANKA^(Balb/c)^ and NK65^(C57BL/6)^ infections show similarities in disease course as well as in the immune responses they trigger. In contrast ANKA^(Balb/c)^ and ANKA(C57BL/6) differed dramatically in each aspects of the disease (Fig.1-3). To reveal whether these observations can be related to differential parasite gene expression, we performed RNA sequencing on blood obtained from infected mice on day 6 pi. Principle component analysis (PCA) of the RNA sequencing showed overlapping patterns for ANKA^(Balb/c)^ and NK65^(C57BL/6)^ infections whereas ANKA^(C57BL/6)^ clustered at a distinct location (Fig.4a,b). Similarly, three way comparative analysis of infected groups showed similar parasite gene expression patterns with subtle differences between ANKA^(Balb/c)^ and NK65^(C57BL/6)^ whereas ANKA^(Balb/c)^ vs ANKA^(C57BL/6)^ and ANKA^(C57BL/6)^ vs NK65^(C57BL/6)^ showed major differences in both the number of differentially expressed genes and their expression patterns (Fig.4c,d) (Supplementary Data 1, Supplementary Fig.1b-d). We performed a thorough gene ontology pathway analysis on the transcriptome and found that the biological processes that were differentially affected between ANKA^(C57BL/6)^ and the other groups were mainly related to parasitic motility in host, protein turnover, exit from host cells and parasite metabolism (Fig. 5 a,b). In terms of molecular function, processes related to catalytic activity of various enzymes, oxidoreductase activity, hydrolase and peptidase activity were predominantly different between ANKA^(C57BL/6)^ and the other groups (Fig 5.c,d). Neither biological process focused pathway analysis, nor molecular function focused pathway analysis revealed a significant difference in parasite behavior between ANKA^(Balb/c)^ and NK65^(C57BL/6)^ indicating that these two groups are indeed highly similar. This may explain similarities in disease progression and outcome as observed in Fig 1-3. Nevertheless, 70 genes were revealed to be differentially expressed between ANKA^(Balb/c)^ and NK65^(C57BL/6)^ Majority of these differences (38 of 70) belonged to interspersed repeat (IR) of the *Berghei* IR (BIR) gene family, *P*.*berghei* strain specific nomenclature of genes of *Plasmodium* IR family (PIR) family (Supplemental Data). PIR, the largest gene family in *Plasmodium*, is composed of genes that are thought to be encoding proteins expressed on iRBCs and are responsible for antigenic variation of parasite^34^. Among the remaining differentially expressed genes, 4 of 70 belong to fam-a, and 6 of 70 belong to fam-b gene families which, like PIRs, are thought to code for iRBC surface proteins that play role in immune evasion, antigenic variation and entry of parasite into host RBCs^27, 35, 36^.

**Figure 4.**
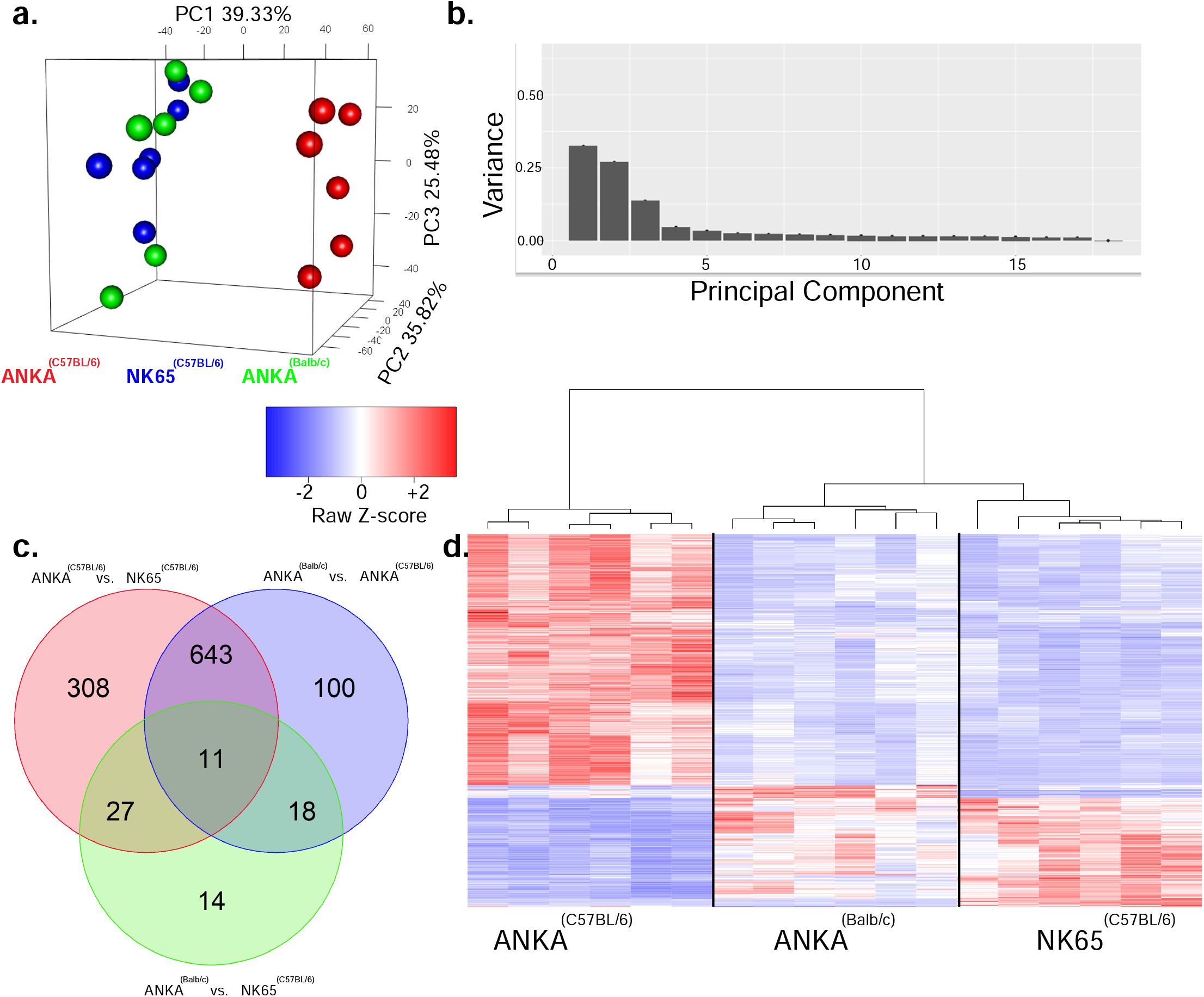
Host determines parasite gene expression. **a-h)** Mice were infected as outlined in Figure 1 and blood samples were taken at day 6 post infection. Changes in parasite gene expression were analyzed by RNA seq carried on 6 blood samples per each group. 3D principal component analysis **(a)** and scree plot **(b)** are shown. Differentially expressing parasite genes (DEG)s between groups are shown as numbers of DEGs in Venn diagram **(c)** and as heat map **(d)**.

**Figure 5.**
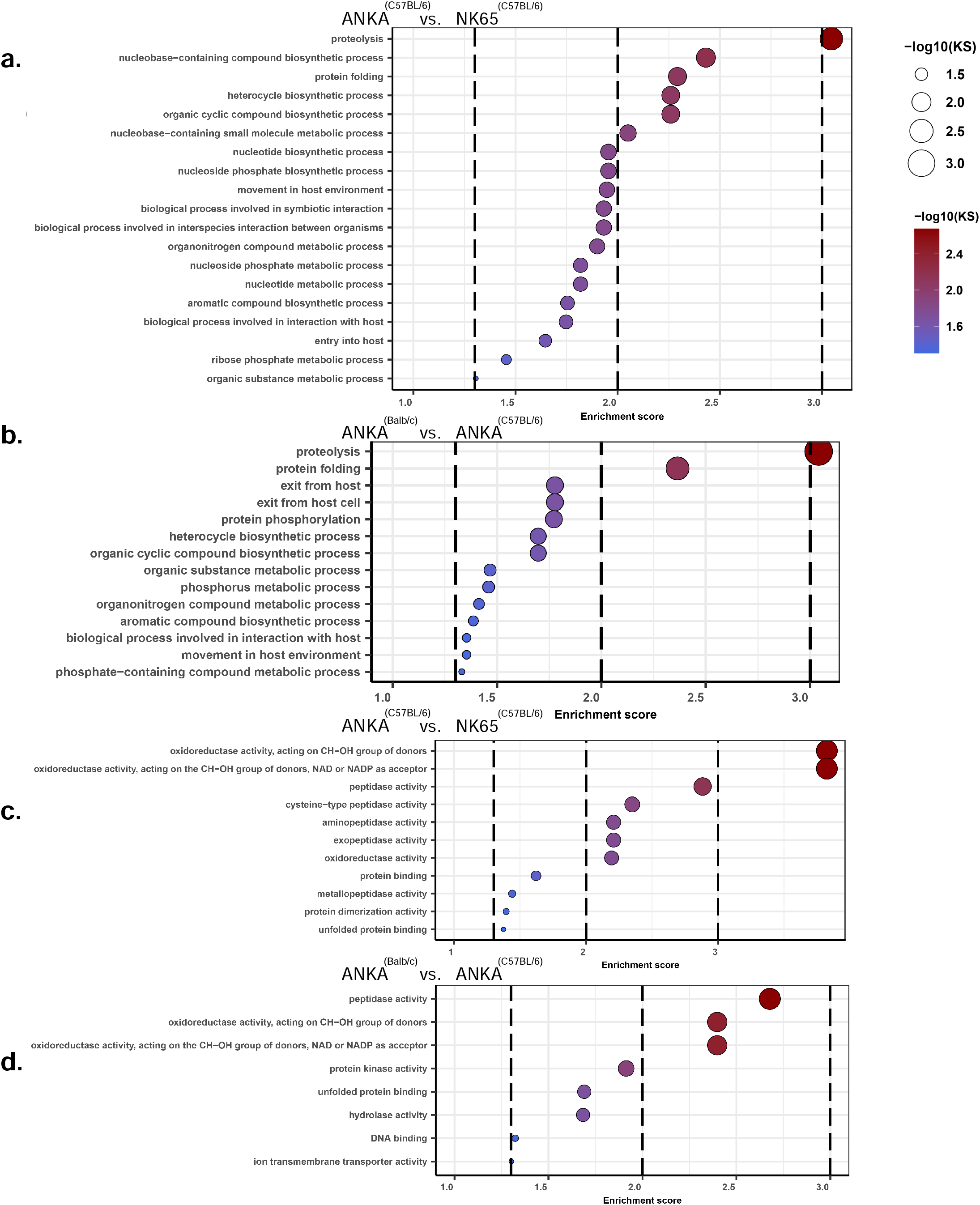
Gene ontology pathway analyses reveals parasite biological pathways that are influenced by host. **a-d)** Gene ontology analyses were performed on the of the RNA seq experiment data shown on Fig.4 focusing on differentially regulated biological processes **(a, b)** and molecular functions **(c, d)**. Dashed vertical cut off lines are drawn at p= 0.05; p=0.01 and p=0.001 (Kolmogorov-Smirnov p-value).

## Discussion

CM is an important public health problem especially among young children living in sub-Saharan Africa. Despite ever growing scientific research it remains unclear why, only a small proportion, approximately 1%, of uncomplicated malaria cases progresses to CM and what determines survival. Animal models have widely been used to overcome the limitations of studying CM using human samples. Studies on ECM in mice have revealed that complex interactions between *Plasmodium* parasites and host immune system play role in the disease pathogenesis. For instance, while CD8^+^ T cells are essential for clearing parasite^37, 38^, their accumulation in brain is linked to development of ECM^19, 39^. Similarly, multiple soluble mediators and humoral factors have been linked to both anti-malarial host defense and triggering brain pathogenesis^28, 40-42^. Separate studies showed that very closely related *Plasmodium* strains such as *Pb* NK65 and *Pb* ANKA can have dramatic differences in their ability to trigger ECM^26^. Therefore, it is apparent that CM is mediated by factors related to both parasite and host.

Here, we provide evidence that in ECM resistant Balb/c host, the ECM causing *Pb* ANKA strain behaves more similarly to the non-ECM causing *Pb* NK65 strain in susceptible C57BL/6 host than *Pb* ANKA infections in C57BL/6 host. We showed that ANKA^(Balb/c)^ and NK65^(C57BL/6)^ infections have overlapping disease courses, both having no symptoms of ECM, and mortality occurring in 21 days due to severe anemia. We also show that unlike ANKA^(C57BL/6)^ which failed to trigger a strong immune response, ANKA^(Balb/c)^ and NK65^(C57BL/6)^ infections induce prominent B and T cell activation/ differentiation. *Plasmodium berghei* infection model is fatal regardless of the host and parasite strain. However, our data showing strong systemic immune response against pathogen ANKA^(Balb/c)^ and NK65^(C57BL/6)^ may, in part, be responsible for preventing rapid progression of the disease to ECM.

In terms of how hosts modulate disease progression, we revealed that, hosts have a strong influence on parasite gene expression turning an ECM causing parasite into a non-ECM causing one. Our analysis on differentially expressed genes between infection groups showed ANKA^(Balb/c)^ and NK65^(C57BL/6)^ differed from ANKA^(C57BL/6)^ with 989 and 772 genes respectively while there were only 70 genes differentially expressed between ANKA^(Balb/c)^ and NK65^(C57BL/6)^. These data explain why ANKA^(Balb/c)^ and NK65^(C57BL/6)^ had so many similarities during the disease progression and why ANKA^(C57BL/6)^ behaved differently. With in-depth pathway analyses, we further revealed that major differences in parasite gene expression between non-ECM and ECM causing conditions cluster around biological processes related to parasite metabolism, protein turnover and parasite’s motility within the host. Therefore, it is apparent that Balb/c host modulates *Pb* ANKA gene expression related to these processes in a way that it is comparable to gene expression in *Pb* NK65 and thus *Pb* ANKA is no longer able to trigger ECM. Likewise, the same set of transcriptional alterations are responsible for longer disease course characterized by stronger immune response seen in both ANKA^(Balb/c)^ and NK65^(C57BL/6)^. Despite remarkable similarity we observed in parasite gene expression between ANKA^(Balb/c)^ and NK65^(C57BL/6)^, there are still 70 differentially expressed genes. Among these genes 48 of 70 belong to BIR, fam-a or fam-b gene families which are collectively thought to regulate antigenic variation and host pathogen interaction. These genes were recently shown to express at different times during blood stage of the parasite and may account for slight variation in the distribution of ring, trophozoite and schizont forms^27^. Altogether, these changes might be a result of adaptation of parasites to different characteristics of C57BL/6 and Balb/c hosts and may account for slight differences between disease courses.

In conclusion, through our comprehensive multidimensional comparative analyses of ANKA^(C57BL/6)^, ANKA^(Balb/c)^ and NK65^(C57BL/6)^ infections, we revealed, in great detail, how host can modulate parasite behavior, dictating the outcome of malaria disease. Our observations in mouse studies can also be linked to human CM in terms of showing how important host genetic background is in determining whether or not blood stage malaria progresses to cerebral disease. Our studies highlight the complex nature of ECM with hundreds of parasite genes altering their expression between ECM causing and non-ECM causing conditions. Nevertheless, the genes and pathways we identify here will facilitate further research on the pathogenesis of the disease.

## Methods

### Mice and Parasites

For all experiments female mice aged 7-8 weeks old were used. Balb/c (Stock no: 000651) and C57BL/6 (stock no: 000664) were purchased from Jackson Laboratories and were maintained in National Institute of Allergy and Infectious Diseases animal facility according to Animal Care and Use Committee Standards. For parasite infections *Plasmodium berghei* NK65 (NYU strain) and *Plasmodium berghei* ANKA strains were used.

### Ethical Statement

Live animal experiments were carried out according to NIAID approved Institutional Animal Care and Use Committee (IACUC) protocol LIG-2E and the Ohio State University approved IACUC protocol 2022A00000061.

### Reagents

Following anti mouse antibodies and other fluorescent markers were used for parasitemia checks, flow cytometry analysis and immunohistochemistry: CXCR5 (BV421-Biolegend), CD4 (BV605-Biolegend), CD8 (Percp Cy 5.5-Biolegend). PD1 (PE-Biolegend), ICOS (AF647-Biolegend), CD44 (AF700-Biolegend), GL7 (e660-Thermofisher), CD95 (AF488-Biolegend), CD19 (BV650-Biolegend), CD138 (PE-Biolegend), Live/Dead near IR (Thermofisher), CD16/32 (Fc block-Biolegend), B220 (AF700-Biolegend), CD45 (APC-Biolegend), Ter119 (APC Cy7-Biolegend), PNA (biotin-Vector Labs), IgM-(eBioscience).

### Infection of animals with parasitized RBCs

Both *Pb* ANKA and *Pb* NK65 parasites stocks were originally grown in C57BL/6 mice. To avoid non-parasite specific immune response against infected C57BL/6 RBCs in Balb/c animals, initial *Pb* ANKA stock was divided into two and injected into both C57BL/6 and Balb/c donor animals as outlined in (Supplementary Fig. 1a). These parasites were passaged three times in respective mouse strains in order to fully adapt the parasite into the host. This strategy also ensured using comparable passage numbers for all animals. Following this adaptation, blood drawn from the third donor, once parasitemia reached to 5%-10% range, was diluted in phosphate buffered saline and injected into experimental animals intraperitoneally 10^6^ iRBCs per mouse.

### Monitoring disease progression in infected animals

Progression of blood stage parasite was monitored by routine parasitemia checks using blood smear and flow cytometry based methods described in detail earlier^43, 44^. Hemoglobin levels were measured using HemoCue Hb201 analyzer. Disease related worsening of clinical symptoms were analyzed using a 10 point clinical scoring system as described earlier^26^. A total clinical score of at least 6 and hemoglobin levels of 2.5 g/dl or below were determined as end-point criteria. Animals meeting one or both criteria were euthanized.

### Flow Cytometry and Tissue section analyses

Spleens, harvested from infected animals were meshed, RBCs were removed using ACK buffer (Lonza). Cells were stained with fluorochrome conjugated antibodies and analyzed in BD X20 flow cytometer. Flow cytometry data was analyzed in FlowJo software vol.10. For histological analyses, Whole brain and spleens were harvested, fixed in 10% buffered formalin and embedded in paraffin. Brain sections were stained with hematoxylin and eosin (H&E). Spleen sections were stained with PNA and IgM and visualized using the strategy outlined in^42^.

### RNA sequencing and data analysis

Mouse cheek blood samples were mixed with TRIzol LS (Thermofisher) in 1:3 ratio and RNA was isolated according to manufacturer’s guidelines. Using a drop of the same blood sample, blood smears were made and distribution of trophozoite, schizont and ring forms of blood stage parasite were quantified. Samples with comparable distributions of these forms between infection groups were selected for next generation sequencing (NGS). Six samples per condition containing 4 μg of total RNA were used for TruSeq mRNA library preparation (Illumina) after being treated with Globin-Zero Gold rRNA Removal Kit (Illumina). Bioanalzyzer DNA 1000 chip was used to fragment-size libraries. Kapa Library Quant kit (Illumina) and Universal qPCR mix (Roche) were used to quantitate libraries to facilitate the generation of a normalized, 2 nM multiplexed pool. This pool was clustered around two RAPID flow cell lanes at 10 pM concentrations. Paired end sequencing was carried out on Illumina HiSeq (100 cycles from fragment ends plus 7 more cycles to sequence the index).

Cutadapt v1.12^45^ was used to remove adapter sequences from RAW NGS data. Reads were then trimmed and filtered (35bp or longer) using FastX Tool Kit v0.0.14 (Hannon Lab, Cold Spring Harbor Laboratory). Hisat2 v2.0.5^46^ set to report only matched pairs was used for mapping. Deseq2 (10.18129/B9.bioc.DESeq210.18129/B9.bioc.DESeq210.18129/B9.bioc.DESeq2) was then used to generate final transcripts based on combined replicates and differentials for each comparison. The sequencing data have been deposited in NCBI’s Gene Expression Omnibus and are accessible through GEO Series accession number GSE215359 (https://www.ncbi.nlm.nih.gov/geo/query/acc.cgi?acc=GSE215359).

### Gene Ontology (GO) Term Enrichment

GO terms annotation file for *Plasmodium berghei* was acquired from PlasmoDB data base (June 2022 release). Enrichment was performed as explained earlier^47^ using R software version 4.2.1 with topGO package^48^.

### Data Analysis

Heatmaps were created using R software version 4.2.1 with heatmap2 package. Statistical analyses were carried out using GraphPad Prism (vol. 7) software.

## Acknowledgements

This study was funded by intramural research grants from National Institute of Allergy and Infectious Diseases allocated to Susan K Pierce and research funds from the Ohio State University College of Medicine allocated to Munir Akkaya. Authors declare no conflicts of interest.

## Author Contributions

MA designed the project. MA, MP, CKC, AS, BPT, MVP, CQ carried out the experiments. MA, CC, MVP, DS, PLC, SMG, SKP, LHM analyzed the data. MA wrote the manuscript. SKP edited the manuscript. MA, SKP secured funding.

## Figure Legends

**Suplementary Figure 1:**
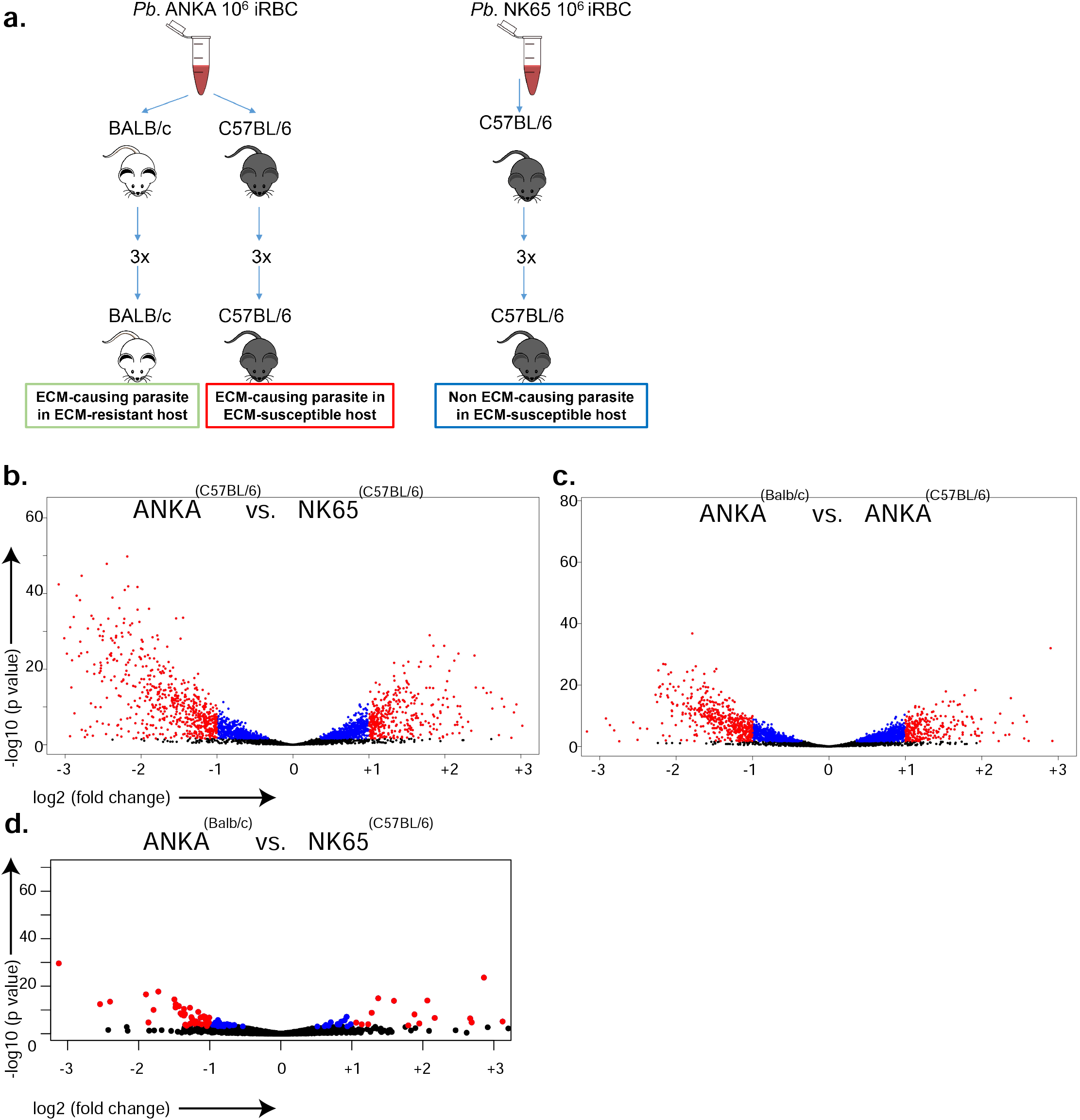
**a)** Outline of the experimental design used for parasite infections. 3X stands for three times passaging the parasite in relevant host before experiment. **b**,**d)** Volcano plots showing the distribution of differentially expressed genes between groups. Genes in red are significant hits.

## References

1. WHO. World Malaria Report. Geneva: World Health Organization, 2021 CC BY-NC-SA 3.0 IGO.

2. Renia L, Grau GE, Wassmer SC. CD8+ T cells and human cerebral malaria: a shifting episteme. J Clin Invest. 2020;130(3):1109–11. Epub 2020/02/18. doi: 10.1172/JCI135510. PubMed PMID: 32065593; PMCID: PMC7269553.

3. Riggle BA, Miller LH, Pierce SK. Desperately Seeking Therapies for Cerebral Malaria. J Immunol. 2020;204(2):327–34. Epub 2020/01/08. doi: 10.4049/jimmunol.1900829. PubMed PMID: 31907275; PMCID: PMC6951433.

4. Idro R, Marsh K, John CC, Newton CR. Cerebral malaria: mechanisms of brain injury and strategies for improved neurocognitive outcome. Pediatr Res. 2010;68(4):267–74. doi: 10.1203/00006450-201011001-0052410.1203/PDR.0b013e3181eee738. PubMed PMID: 20606600; PMCID: PMC3056312.

5. Idro R, Kakooza-Mwesige A, Balyejjussa S, Mirembe G, Mugasha C, Tugumisirize J, Byarugaba J. Severe neurological sequelae and behaviour problems after cerebral malaria in Ugandan children. BMC Res Notes. 2010;3:104. Epub 20100416. doi: 10.1186/1756-0500-3-104. PubMed PMID: 20398391; PMCID: PMC2861066.

6. Holmberg D, Franzen-Rohl E, Idro R, Opoka RO, Bangirana P, Sellgren CM, Wickstrom R, Farnert A, Schwieler L, Engberg G, John CC. Cerebrospinal fluid kynurenine and kynurenic acid concentrations are associated with coma duration and long-term neurocognitive impairment in Ugandan children with cerebral malaria. Malar J. 2017;16(1):303. Epub 20170728. doi: 10.1186/s12936-017-1954-1. PubMed PMID: 28754152; PMCID: PMC5534063.

7. Riggle BA, Miller LH, Pierce SK. Do we know enough to find an adjunctive therapy for cerebral malaria in African children? F1000Res. 2017;6:2039. Epub 2017/12/19. doi: 10.12688/f1000research.12401.1. PubMed PMID: 29250318; PMCID: PMC5701444.

8. MacPherson GG, Warrell MJ, White NJ, Looareesuwan S, Warrell DA. Human cerebral malaria. A quantitative ultrastructural analysis of parasitized erythrocyte sequestration. Am J Pathol. 1985;119(3):385–401. PubMed PMID: 3893148; PMCID: PMC1888001.

9. Nickerson JP, Tong KA, Raghavan R. Imaging cerebral malaria with a susceptibility-weighted MR sequence. AJNR Am J Neuroradiol. 2009;30(6):e85–6. Epub 20090325. doi: 10.3174/ajnr.A1568. PubMed PMID: 19321625; PMCID: PMC7051359.

10. Riggle BA, Manglani M, Maric D, Johnson KR, Lee MH, Neto OLA, Taylor TE, Seydel KB, Nath A, Miller LH, McGavern DB, Pierce SK. CD8+ T cells target cerebrovasculature in children with cerebral malaria. J Clin Invest. 2020;130(3):1128–38. doi: 10.1172/JCI133474. PubMed PMID: 31821175; PMCID: PMC7269583.

11. Seydel KB, Kampondeni SD, Valim C, Potchen MJ, Milner DA, Muwalo FW, Birbeck GL, Bradley WG, Fox LL, Glover SJ, Hammond CA, Heyderman RS, Chilingulo CA, Molyneux ME, Taylor TE. Brain swelling and death in children with cerebral malaria. N Engl J Med. 2015;372(12):1126–37. doi: 10.1056/NEJMoa1400116. PubMed PMID: 25785970; PMCID: PMC4450675.

12. Newton CR, Hien TT, White N. Cerebral malaria. J Neurol Neurosurg Psychiatry. 2000;69(4):433–41. doi: 10.1136/jnnp.69.4.433. PubMed PMID: 10990500; PMCID: PMC1737146.

13. Palmiere C, Jaton K, Lobrinus A, Schrag B, Greub G. Post-mortem diagnosis of malaria. New Microbes New Infect. 2014;2(5):154–5. Epub 20140715. doi: 10.1002/nmi2.52. PubMed PMID: 25356366; PMCID: PMC4184481.

14. Idro R, Jenkins NE, Newton CR. Pathogenesis, clinical features, and neurological outcome of cerebral malaria. Lancet Neurol. 2005;4(12):827–40. doi: 10.1016/S1474-4422(05)70247-7. PubMed PMID: 16297841.

15. Huang BW, Pearman E, Kim CC. Mouse Models of Uncomplicated and Fatal Malaria. Bio Protoc. 2015;5(13). doi: 10.21769/bioprotoc.1514. PubMed PMID: 26236758; PMCID: PMC4520541.

16. Ghazanfari N, Mueller SN, Heath WR. Cerebral Malaria in Mouse and Man. Front Immunol. 2018;9:2016. Epub 2018/09/27. doi: 10.3389/fimmu.2018.02016. PubMed PMID: 30250468; PMCID: PMC6139318.

17. Riggle BA, Sinharay S, Schreiber-Stainthorp W, Munasinghe JP, Maric D, Prchalova E, Slusher BS, Powell JD, Miller LH, Pierce SK, Hammoud DA. MRI demonstrates glutamine antagonist-mediated reversal of cerebral malaria pathology in mice. Proc Natl Acad Sci U S A. 2018;115(51):E12024–E33. Epub 2018/12/06. doi: 10.1073/pnas.1812909115. PubMed PMID: 30514812; PMCID: PMC6304986.

18. Randall LM, Amante FH, McSweeney KA, Zhou Y, Stanley AC, Haque A, Jones MK, Hill GR, Boyle GM, Engwerda CR. Common strategies to prevent and modulate experimental cerebral malaria in mouse strains with different susceptibilities. Infect Immun. 2008;76(7):3312–20. Epub 20080512. doi: 10.1128/IAI.01475-07. PubMed PMID: 18474652; PMCID: PMC2446712.

19. Swanson PA, 2nd, Hart GT, Russo MV, Nayak D, Yazew T, Pena M, Khan SM, Janse CJ, Pierce SK, McGavern DB. CD8+ T Cells Induce Fatal Brainstem Pathology during Cerebral Malaria via Luminal Antigen-Specific Engagement of Brain Vasculature. PLoS Pathog. 2016;12(12):e1006022. Epub 2016/12/03. doi: 10.1371/journal.ppat.1006022. PubMed PMID: 27907215; PMCID: PMC5131904.

20. Gordon EB, Hart GT, Tran TM, Waisberg M, Akkaya M, Kim AS, Hamilton SE, Pena M, Yazew T, Qi CF, Lee CF, Lo YC, Miller LH, Powell JD, Pierce SK. Targeting glutamine metabolism rescues mice from late-stage cerebral malaria. Proc Natl Acad Sci U S A. 2015;112(42):13075–80. doi: 10.1073/pnas.1516544112. PubMed PMID: 26438846; PMCID: PMC4620853.

21. Grau GE, Piguet PF, Engers HD, Louis JA, Vassalli P, Lambert PH. L3T4+ T lymphocytes play a major role in the pathogenesis of murine cerebral malaria. J Immunol. 1986;137(7):2348–54. PubMed PMID: 3093572.

22. Hanum PS, Hayano M, Kojima S. Cytokine and chemokine responses in a cerebral malaria-susceptible or -resistant strain of mice to Plasmodium berghei ANKA infection: early chemokine expression in the brain. Int Immunol. 2003;15(5):633–40. doi: 10.1093/intimm/dxg065. PubMed PMID: 12697663.

23. Mackey LJ, Hochmann A, June CH, Contreras CE, Lambert PH. Immunopathological aspects of Plasmodium berghei infection in five strains of mice. II. Immunopathology of cerebral and other tissue lesions during the infection. Clin Exp Immunol. 1980;42(3):412–20. PubMed PMID: 7011607; PMCID: PMC1537166.

24. Rest JR. Cerebral malaria in inbred mice. I. A new model and its pathology. Trans R Soc Trop Med Hyg. 1982;76(3):410–5. doi: 10.1016/0035-9203(82)90203-6. PubMed PMID: 7051459.

25. Neill AL, Chan-Ling T, Hunt NH. Comparisons between microvascular changes in cerebral and non-cerebral malaria in mice, using the retinal whole-mount technique. Parasitology. 1993;107 (Pt 5):477–87. doi: 10.1017/s0031182000068050. PubMed PMID: 8295787.

26. Akkaya M, Bansal A, Sheehan PW, Pena M, Cimperman CK, Qi CF, Yazew T, Otto TD, Billker O, Miller LH, Pierce SK. Testing the impact of a single nucleotide polymorphism in a Plasmodium berghei ApiAP2 transcription factor on experimental cerebral malaria in mice. Sci Rep. 2020;10(1):13630. Epub 2020/08/14. doi: 10.1038/s41598-020-70617-7. PubMed PMID: 32788672; PMCID: PMC7424516.

27. Otto TD, Bohme U, Jackson AP, Hunt M, Franke-Fayard B, Hoeijmakers WA, Religa AA, Robertson L, Sanders M, Ogun SA, Cunningham D, Erhart A, Billker O, Khan SM, Stunnenberg HG, Langhorne J, Holder AA, Waters AP, Newbold CI, Pain A, Berriman M, Janse CJ. A comprehensive evaluation of rodent malaria parasite genomes and gene expression. BMC Biol. 2014;12:86. doi: 10.1186/s12915-014-0086-0. PubMed PMID: 25359557; PMCID: PMC4242472.

28. Akkaya M, Bansal A, Sheehan PW, Pena M, Molina-Cruz A, Orchard LM, Cimperman CK, Qi CF, Ross P, Yazew T, Sturdevant D, Anzick SL, Thiruvengadam G, Otto TD, Billker O, Llinas M, Miller LH, Pierce SK. A single-nucleotide polymorphism in a Plasmodium berghei ApiAP2 transcription factor alters the development of host immunity. Sci Adv. 2020;6(6):eaaw6957. Epub 2020/02/23. doi: 10.1126/sciadv.aaw6957. PubMed PMID: 32076635; PMCID: PMC7002124.

29. Zhao H, Aoshi T, Kawai S, Mori Y, Konishi A, Ozkan M, Fujita Y, Haseda Y, Shimizu M, Kohyama M, Kobiyama K, Eto K, Nabekura J, Horii T, Ishino T, Yuda M, Hemmi H, Kaisho T, Akira S, Kinoshita M, Tohyama K, Yoshioka Y, Ishii KJ, Coban C. Olfactory plays a key role in spatiotemporal pathogenesis of cerebral malaria. Cell Host Microbe. 2014;15(5):551–63. doi: 10.1016/j.chom.2014.04.008. PubMed PMID: 24832450.

30. Yanez DM, Manning DD, Cooley AJ, Weidanz WP, van der Heyde HC. Participation of lymphocyte subpopulations in the pathogenesis of experimental murine cerebral malaria. J Immunol. 1996;157(4):1620–4. Epub 1996/08/15. PubMed PMID: 8759747.

31. de Kossodo S, Grau GE. Profiles of cytokine production in relation with susceptibility to cerebral malaria. J Immunol. 1993;151(9):4811–20. PubMed PMID: 8409439.

32. Griffith JW, O’Connor C, Bernard K, Town T, Goldstein DR, Bucala R. Toll-like receptor modulation of murine cerebral malaria is dependent on the genetic background of the host. J Infect Dis. 2007;196(10):1553–64. Epub 20071031. doi: 10.1086/522865. PubMed PMID: 18008236.

33. Shibui A, Hozumi N, Shiraishi C, Sato Y, Iida H, Sugano S, Watanabe J. CD4(+) T cell response in early erythrocytic stage malaria: Plasmodium berghei infection in BALB/c and C57BL/6 mice. Parasitol Res. 2009;105(1):281–6. Epub 20090408. doi: 10.1007/s00436-009-1435-8. PubMed PMID: 19352703.

34. Cunningham D, Lawton J, Jarra W, Preiser P, Langhorne J. The pir multigene family of Plasmodium: antigenic variation and beyond. Mol Biochem Parasitol. 2010;170(2):65–73. doi: 10.1016/j.molbiopara.2009.12.010. PubMed PMID: 20045030.

35. Alam MS, Zeeshan M, Rathore S, Sharma YD. Multiple Plasmodium vivax proteins of Pvfam-a family interact with human erythrocyte receptor Band 3 and have a role in red cell invasion. Biochem Biophys Res Commun. 2016;478(3):1211–6. Epub 20160818. doi: 10.1016/j.bbrc.2016.08.096. PubMed PMID: 27545606.

36. Zeeshan M, Tyagi RK, Tyagi K, Alam MS, Sharma YD. Host-parasite interaction: selective Pv-fam-a family proteins of Plasmodium vivax bind to a restricted number of human erythrocyte receptors. J Infect Dis. 2015;211(7):1111–20. Epub 20141013. doi: 10.1093/infdis/jiu558. PubMed PMID: 25312039.

37. Safeukui I, Gomez ND, Adelani AA, Burte F, Afolabi NK, Akondy R, Velazquez P, Holder A, Tewari R, Buffet P, Brown BJ, Shokunbi WA, Olaleye D, Sodeinde O, Kazura J, Ahmed R, Mohandas N, Fernandez-Reyes D, Haldar K. Malaria induces anemia through CD8+ T cell-dependent parasite clearance and erythrocyte removal in the spleen. mBio. 2015;6(1). Epub 20150120. doi: 10.1128/mBio.02493-14. PubMed PMID: 25604792; PMCID: PMC4324318.

38. Imai T, Ishida H, Suzue K, Taniguchi T, Okada H, Shimokawa C, Hisaeda H. Cytotoxic activities of CD8(+) T cells collaborate with macrophages to protect against blood-stage murine malaria. Elife. 2015;4. Epub 20150311. doi: 10.7554/eLife.04232. PubMed PMID: 25760084; PMCID: PMC4366679.

39. Barrera V, Haley MJ, Strangward P, Attree E, Kamiza S, Seydel KB, Taylor TE, Milner DA, Jr., Craig AG, Couper KN. Comparison of CD8(+) T Cell Accumulation in the Brain During Human and Murine Cerebral Malaria. Front Immunol. 2019;10:1747. Epub 20190724. doi: 10.3389/fimmu.2019.01747. PubMed PMID: 31396236; PMCID: PMC6668485.

40. Angulo I, Fresno M. Cytokines in the pathogenesis of and protection against malaria. Clin Diagn Lab Immunol. 2002;9(6):1145–52. doi: 10.1128/cdli.9.6.1145-1152.2002. PubMed PMID: 12414742; PMCID: PMC130117.

41. Villegas-Mendez A, Strangward P, Shaw TN, Rajkovic I, Tosevski V, Forman R, Muller W, Couper KN. Gamma Interferon Mediates Experimental Cerebral Malaria by Signaling within Both the Hematopoietic and Nonhematopoietic Compartments. Infect Immun. 2017;85(11). Epub 20171018. doi: 10.1128/IAI.01035-16. PubMed PMID: 28874445; PMCID: PMC5649021.

42. Akkaya M, Akkaya B, Sheehan PW, Miozzo P, Pena M, Qi CF, Manzella-Lapeira J, Bolland S, Pierce SK. T cell-dependent antigen adjuvanted with DOTAP-CpG-B but not DOTAP-CpG-A induces robust germinal center responses and high affinity antibodies in mice. Eur J Immunol. 2017;47(11):1890–9. Epub 2017/08/02. doi: 10.1002/eji.201747113. PubMed PMID: 28762497.

43. Gordon EB, Hart GT, Tran TM, Waisberg M, Akkaya M, Skinner J, Zinocker S, Pena M, Yazew T, Qi CF, Miller LH, Pierce SK. Inhibiting the Mammalian target of rapamycin blocks the development of experimental cerebral malaria. MBio. 2015;6(3):e00725. doi: 10.1128/mBio.00725-15. PubMed PMID: 26037126; PMCID: PMC4453009.

44. Malleret B, Claser C, Ong AS, Suwanarusk R, Sriprawat K, Howland SW, Russell B, Nosten F, Renia L. A rapid and robust tri-color flow cytometry assay for monitoring malaria parasite development. Sci Rep. 2011;1:118. Epub 2012/02/23. doi: 10.1038/srep00118. PubMed PMID: 22355635; PMCID: PMC3216599.

45. Martin M. Cutadapt removes adapter sequences from high-throughput sequencing reads. EMBnet J. 2011;17:10–2. doi: https://doi.org/10.14806/ej.17.1.200.

46. Kim D, Langmead B, Salzberg SL. HISAT: a fast spliced aligner with low memory requirements. Nat Methods. 2015;12(4):357–60. Epub 2015/03/10. doi: 10.1038/nmeth.3317. PubMed PMID: 25751142; PMCID: PMC4655817.

47. Bushell E, Gomes AR, Sanderson T, Anar B, Girling G, Herd C, Metcalf T, Modrzynska K, Schwach F, Martin RE, Mather MW, McFadden GI, Parts L, Rutledge GG, Vaidya AB, Wengelnik K, Rayner JC, Billker O. Functional Profiling of a Plasmodium Genome Reveals an Abundance of Essential Genes. Cell. 2017;170(2):260–72 e8. doi: 10.1016/j.cell.2017.06.030. PubMed PMID: 28708996; PMCID: PMC5509546.

48. Alexa A, Rahnenfuhrer J, Lengauer T. Improved scoring of functional groups from gene expression data by decorrelating GO graph structure. Bioinformatics. 2006;22(13):1600–7. Epub 20060410. doi: 10.1093/bioinformatics/btl140. PubMed PMID: 16606683.

